# Active modulation of Hydrogen bonding by sericin enhances cryopreservation outcomes

**DOI:** 10.1101/773721

**Authors:** L. Underwood, J. Solocinski, E. Rosiek, Q. Osgood, N. Chakraborty

**Affiliations:** University of Michigan Dearborn; OtziBio Inc

## Abstract

Cryopreservation of cells without any toxicity concerns is a critical step in ensuring successful clinical translation of cell-based technologies. Mitigating the toxicity concerns related to most of the commonly used cryoprotectants including dimethyl sulfoxide (DMSO) is an active area of research in cryobiology. In recent years use of additives including polymeric proteins such has sericin have been explored as an additive to cryoprotectant formulations. In this study the thermophysical effect of addition of sericin was investigated. The effect of presence of sericin on the H-bonding strength was investigated using Raman microspectroscopy and other thermophysical effects were quantified using differential scanning calorimetry (DSC) techniques. Finally, the prospect of using sericin as an additive to cryoprotectant formulation was investigated by monitoring cellular viability and growth following exposure to cryogenic temperatures in hepatocellular carcinoma cells. Results indicate significant improvement in post-thaw viability when sericin is used as an additive to DMSO based formulations. While use of trehalose as an additive has beneficial effects by itself, combined usage of sericin and trehalose as additives did result in an improved overall long-term growth potential of the cells.

**Statement of Significance:** This study provides for powerful biophysical understanding of how sericin can be used as an additive for cryoprotectant solutions, which allows storage of biologics at low temperatures. It is desirable to replace current components of cryoprotectant formulation (such as DMSO) due to innate toxicity and metabolic derangements to cells. The ability of sericin to improve cryoprotective solutions was mechanistically characterized by Raman microspectroscopy, which allows for molecular level characterization of the nature of H-bonding in aqueous environments in presence of solution components. Thermodynamic analysis of the cryoprotectant solutions containing sericin was undertaken to quantify the relation between solution composition and cryopreservation outcome. This analytical study provides a basis for designing better cryoprotectants with lower thermophysical injury and higher cellular yields.

## Introduction

Cell preservation technologies can play an extremely important role in transitioning cell-based techniques from laboratory to bedside. Such translation of cell-based technologies require development of highly optimized and efficient long-term preservation strategy for cells. In general, long-term preservation of cells and cellular materials is achieved through formation of a glassy matrix [1, 2] at low temperatures in presence of cryoprotective agents (CPAs). This glassy matrix has been hypothesized to reduce molecular mobility and prevent degradative intracellular reactions at low storage temperatures [3]. Dimethyl sulfoxide (DMSO) is one of the most commonly used CPAs in slow cooling rate cryopreservation techniques [4, 5]. However, use of DMSO has been frequently and closely associated with cellular toxicity effects and poor post-thaw performance [6–8].

While there are very few credible alternatives to DMSO, at high concentrations (>10%) DMSO has been shown to be toxic to cells as it is known to take part in formation of lipid membranes pores [9] and other irreversible membrane damage. This characteristic has been well-studied in the context of drug delivery development strategies [10]. Even at low concentrations, exposure to DMSO (<4% v/v) has recently been linked to irreversible cellular damage, as it has been shown to initiate apoptosis in retinal neuronal cells [6]. In another study, exposure to 1% v/v DMSO levels did not produce any evidence of cell death, but there was significant mitochondrial damage due to increased levels of reactive oxygen species (ROS) following 24-hour exposure [11]. This leads to mitochondrial swelling and significant membrane potential impairment [12]. A recent study indicates that exposing human tissue to 0.1% v/v levels of DMSO causes drastic changes to human intercellular processes and the overall epigenetic landscape by impacting DNA methylenation and down regulating microRNAs [13]. All of these studies suggest the need to revisit the optimal use of DMSO in CPA formulations.

Use of additives that can actively modulate the cryopreservation outcome in CPA formulations is a commonly accepted strategy. Several disaccharides, including trehalose and sucrose [14–16], glycerol [15], and proline [17, 18] have been used as additives to DMSO-based CPA formulations. The effect of using additives to DMSO based CPA formulations can vary and range from being biophysical to biochemical in nature. For example, when trehalose is used as an additive, it can play a cryoprotective role by reducing ice crystal size and thereby discourage the formation of harmful ice crystal patterns during cryopreservation. Such an effect on crystal formation was found to have a direct positive impact on cellular viability and post-thaw metabolism following cryopreservation [19]. The study also revealed the delicate balance that is needed to avoid the two principal cryogenic injury paradigms while formulating CPAs [4]. While on one hand addition of an additive can reduce intracellular ice formation (IIF), an increase in concentrations beyond a threshold can lead to increased osmotic stress and decreased viability. Additives may also exert their cytoprotective effect by various biochemical means. During slow-cooling rate (<10°C/min) cryopreservation, cells are generally exposed to hyperosmotic CPA solutions for several minutes before extracellular crystallization occurs [4, 20, 21]. Several osmolytes, such as proline [22] and certain anti-apoptotic agents [23, 24] have been shown to play a biochemical role in avoiding osmotic injury during slow-cooling rate cryopreservation.

In this study, the effect of using the polymeric protein sericin as a non-penetrating CPA was investigated. Sericin is a protein used by bombyx mori (silkworms) in the production of silk. Sericin acts as an adhesive coating to the fibers and has anti-oxidant properties [25]. Clinically, sericin shows promise as a protective molecule against several types of cellular stresses. Multiple studies have reported sericin’s ability to mitigate oxidative stress in various tissue types and proposed use of sericin as a replacement for animal origin serum in cell culture [26, 27]. It has been already been used as a serum replacing agent in CPA formulations for several different types of mammalian cells including human adipose tissue-derived stem cells [28], myeloma cell lines, fibroblasts, keratinocytes, insect cell lines [29], rat insulinoma cell lines, mouse hybridoma cell lines and mesenchymal stem cells [30].

Due to the polymeric nature of the protein, sericin can form extensive hydrogen bonding (H-bonding) in the matrix [31] and thus can play an important role in creating low-molecular mobility environment at low temperatures [32]. H-bonding can be used to impact the nature of the glass formation by modulating the nature of H-bonds formed [33] and the H-bonding strength plays a critical role in this regard. In this study an investigation was undertaken to characterize the effect of sericin as an additive to DMSO based CPA formulation. Special emphasis was given in gaining understanding of the effect of addition of sericin in aqueous environment on H-bonding strength using Raman microspectroscopy. Thermophysical properties of CPA formulations containing different concentrations of sericin were characterized using differential scanning calorimetry and these properties were likened to the cryopreservation outcome of human hepatocellular carcinoma cells (HepG2) in terms of cellular survival and growth.

The studies performed here provide a framework for biophysical criteria required towards development of low-toxicity CPA formulations that can be used in highly optimized long-term preservation techniques under clinical settings.

## Materials and Methods

### Raman microspectroscopic analysis of hydrogen bonding of cryoprotective solutions

The thermo-molecular effect of addition of sericin in the CPA formulation was investigated by quantifying H-bonding characteristics and strengths at different temperatures and concentrations using Raman microspectroscopy. In doing so, special emphasis was given to the OH stretching region that can be used to understand the effect of H-bonding in aqueous solutions. Raman spectral measurements were performed using a customized confocal microscope Raman spectrometer (UHTS 300, WITec Instruments Crop, Germany). A 532-nm solid-state laser system was used for photonic excitation. Spectral signatures were collected using a 10X objective (Zeiss, Thornwood, NY) and an EMCCD camera (Andor Technology, UK). A liquid nitrogen-cooled low temperature stage (FDCS 196, Linkam Scientific Instruments, UK) was used to control sample temperatures. The temperature-controlled stage was mounted on the Raman microscope stage using custom-made stage adaptors.

For each experiment, 300 µL of solution was added to a quartz crucible and placed inside the freezing stage. Samples were initially cooled to 0°C then warmed in 5°C increments and allowed to stabilize at each temperature point until reaching 20°C; a heating/cooling rate of 10°C/min was used. Spectra were gathered at each temperature point using an integration time of 1 s, averaged over 60 accumulations. Following collection, background spectra was subtracted, and cosmic ray interference were removed. Spectral peaks were deconvoluted and analyzed using Origin Pro 2018 (OriginLab, Northampton, MA).

A customized chemometric deconvolution algorithm based on Fast Fourier Transform (FFT) of Raman signal is used to decompose the OH stretching regions of the CPA formulations [34, 35]. The deconvolution is computed using the formulation

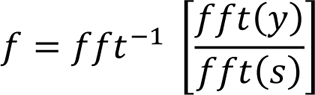

where *y* is the known response of the signal *s*. While several different peaks can be identified in the OH stretching region (Fig. 1A) that are related to the physical state of the water and H-bonding, two principals peaks related to symmetric (∼3200 cm^-1^) and asymmetric (∼3415 cm^-1^) vibrations were considered here for the analysis related to H-bonding. The higher-frequency asymmetric spectral component is known to be related to the water molecules bound by incompletely formed H-bonding [36]. Whereas, the lower-frequency symmetric component corresponds to the molecules with complete tetrahedral H-bonded structure [37, 38].

**Figure 1:**
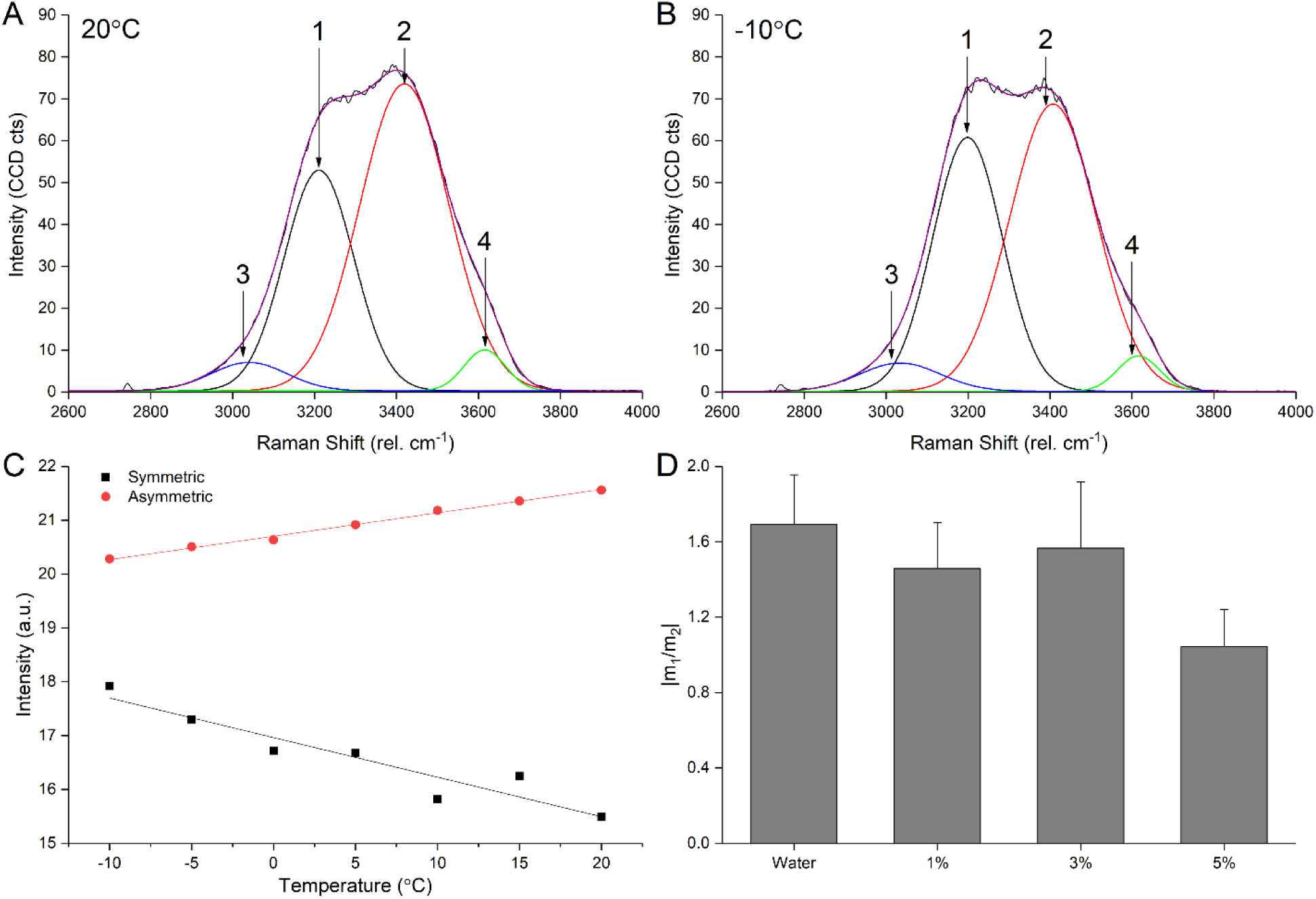
Raman spectroscopy of binary water-sericin solutions A) OH-stretching band (∼2900-3700 cm^-1^) of pure water at −10°C. Spectrum is deconvoluted into four primary bands and the reconstructed spectrum is superimposed on the original to show agreement of fit. Arrows 1 and 2 indicate symmetric and asymmetric peaks respectively. B) OH-stretching band of pure water at 20°C. C) Symmetric and asymmetric peak intensities are plotted at the corresponding temperatures at which Raman scans were acquired for pure water. Linear fits are calculated for both sets of data with R^2^ ≥ 0.9 (sym) and 0.995 (asym). D) The ratio of slopes, m_1_ to m_2_, was calculated for different sericin concentrations in water. Error bars represent SEM of slope fit.

The experimentally obtained spectral intensity of the symmetric and asymmetric peaks in the OH stretching region were then used to estimate the enthalpy and entropy of the formation of hydrogen bonds in aqueous CPA formulations containing sericin as additives. Van’t Hoff equation was used to relate enthalpy change and the equilibrium constant of reaction for CPA solutions [39, 40]. A van’t Hoff plot (Fig. 2A) was constructed as the linear dependence of ln(k) against the inverse of the temperature (T). The enthalpy (ΔH) of bonding is was expressed as a product of the slope of the Van’t Hoff plot and universal gas constant. The equilibrium constant (k) in this case is also equal to the ratio of the intensities of these individual peaks resolved in Raman spectrum at different temperatures (T). [41]

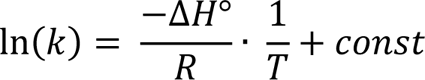

**Figure 2:**
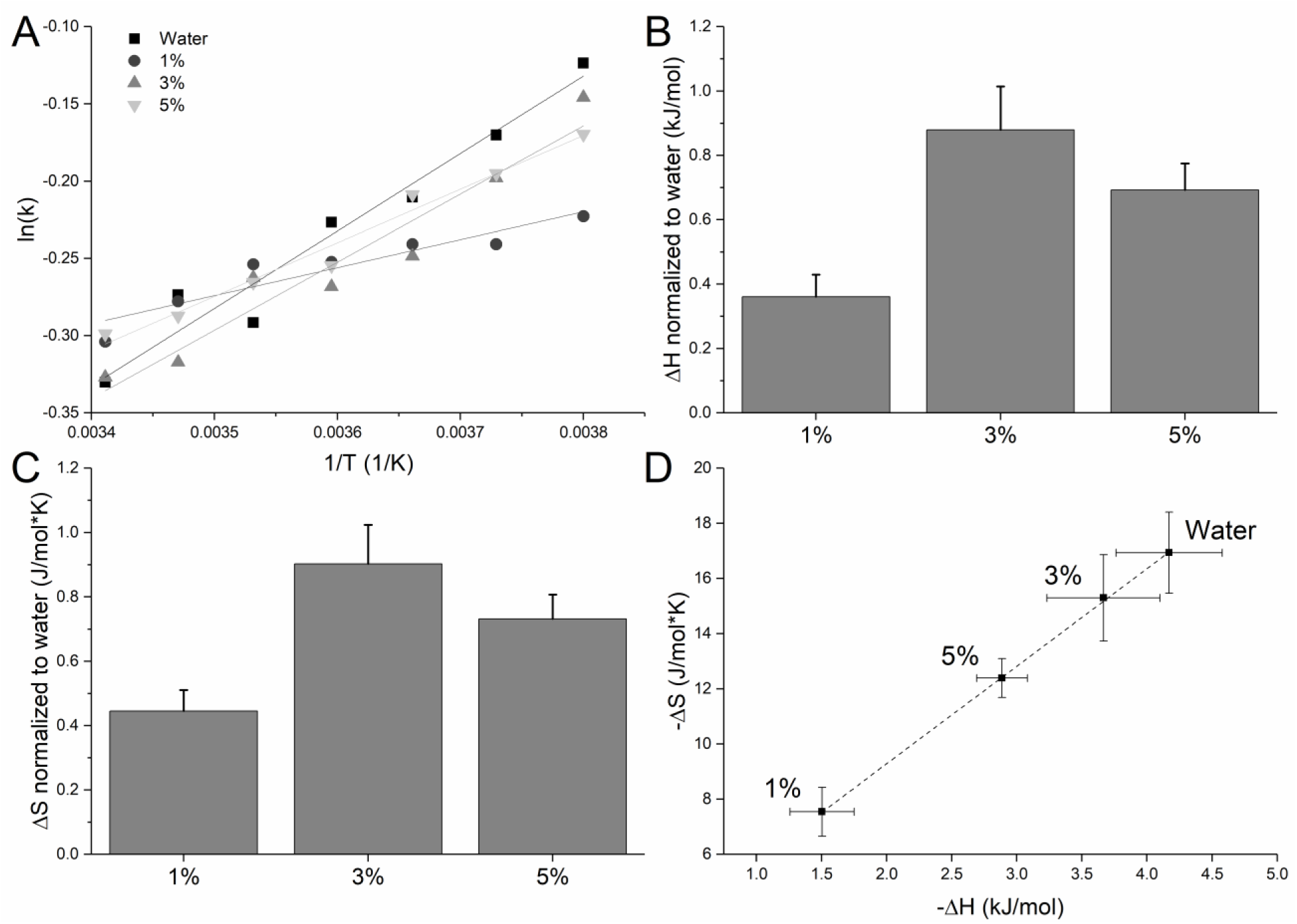
Raman spectroscopic thermodynamic analysis: A) Van’t Hoff plots for water-sericin solutions, k represents ratio between symmetric and asymmetric intensity. B) Change in the enthalpy of water-sericin binary solutions based on Van’t Hoff plots, normalized to pure water. *p < 0.05. C) Change in the entropy of water-sericin binary solutions normalized to pure water. *p < 0.05. D) Linear fit of change in enthalpy against change in entropy for water-sericin binary solutions. Error bars show ±SEM, n=3

Here R is the universal gas constant, and ΔH° is the change in enthalpy during the formation of one mole of hydrogen bonded molecules from nonbonded ones under standard conditions of 298 K and 1 atm. The change in enthalpy (Fig. 2B) and entropy (Fig. 2C) was used to quantify the change in characteristics of H-bonding strength in aqueous solutions having varying concentrations of sericin.

In addition to the H-bonding characterization using intensities of the symmetric and asymmetric peaks in the OH stretching region of the Raman spectra, change of peak position (peak shift) was also used to analyze energetics related to H-bonding. The trend in peak-shift characteristics were quantified by comparing the change in peak-center per unit temperature for both symmetric (n_1_) and asymmetric peaks (n_2_, as seen in Fig. 3A and B). This analysis indicates the variation in H-bonding energetics at different temperatures in presence of sericin.

**Figure 3:**
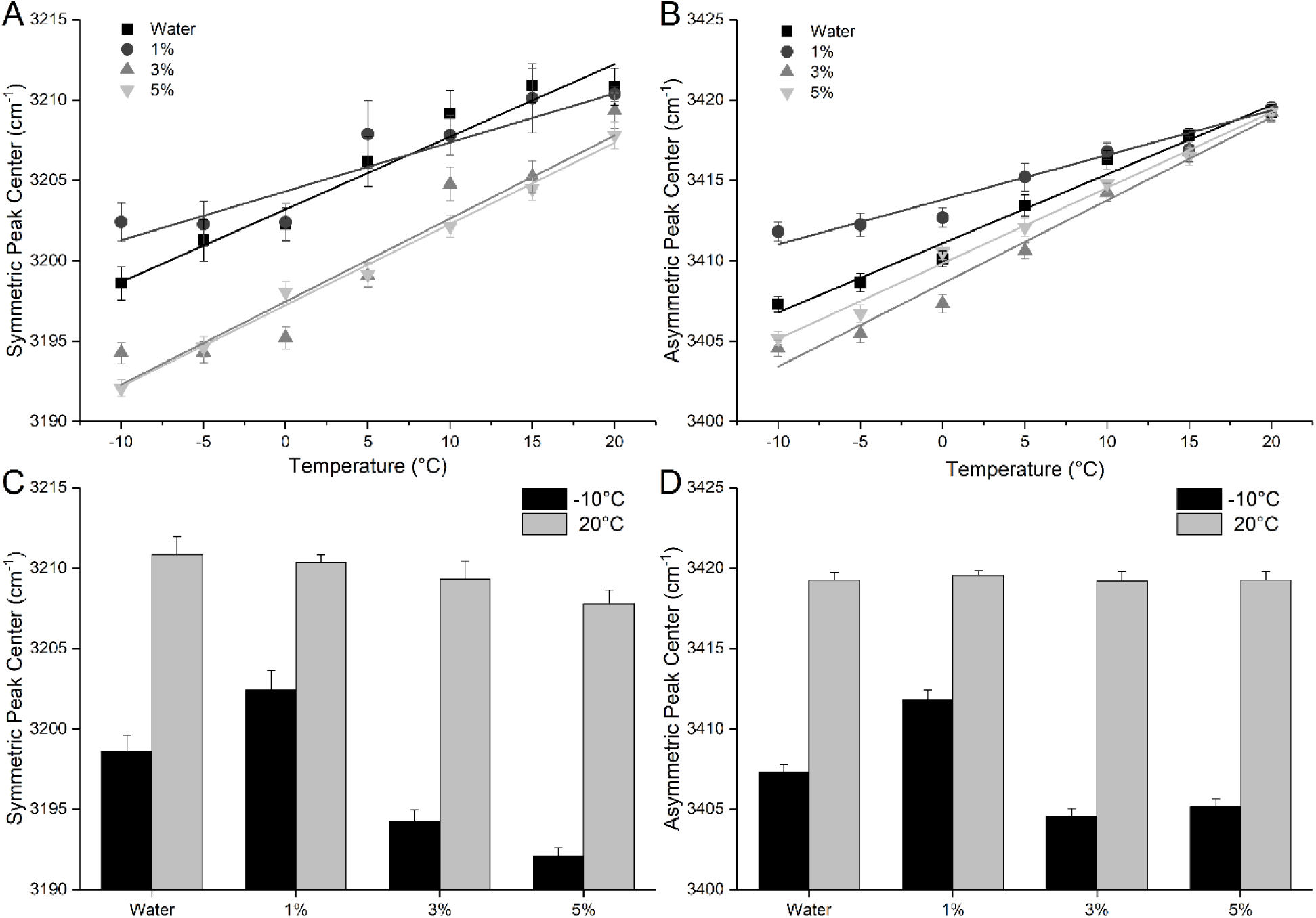
Peak center shift of OH stretching bands A) Symmetric peak center is plotted for water-sericin solutions at temperatures ranging from −10°C to 20°C. A noticeable difference in slope can be observed for the 1% sericin solution. B) Asymmetric peak center plotted for the same solutions and temperatures as (A). As temperature increases, peak center converges asymmetric peak center converges at 3417 cm^-1^. This indicates equal asymmetric hydrogen bonding at higher temperature. C) Symmetric peak center values for various water-sericin solutions at −10°C and 20°C. At low temperature, there are significant differences between the peak center values of the solutions. There is a slight trend towards lower frequency peak centers as sericin increases at higher temperature. D) Asymmetric peak center values for water-sericin solutions. At low temperature, peak center shift follows a similar trend as the symmetric.

**Figure 4:**
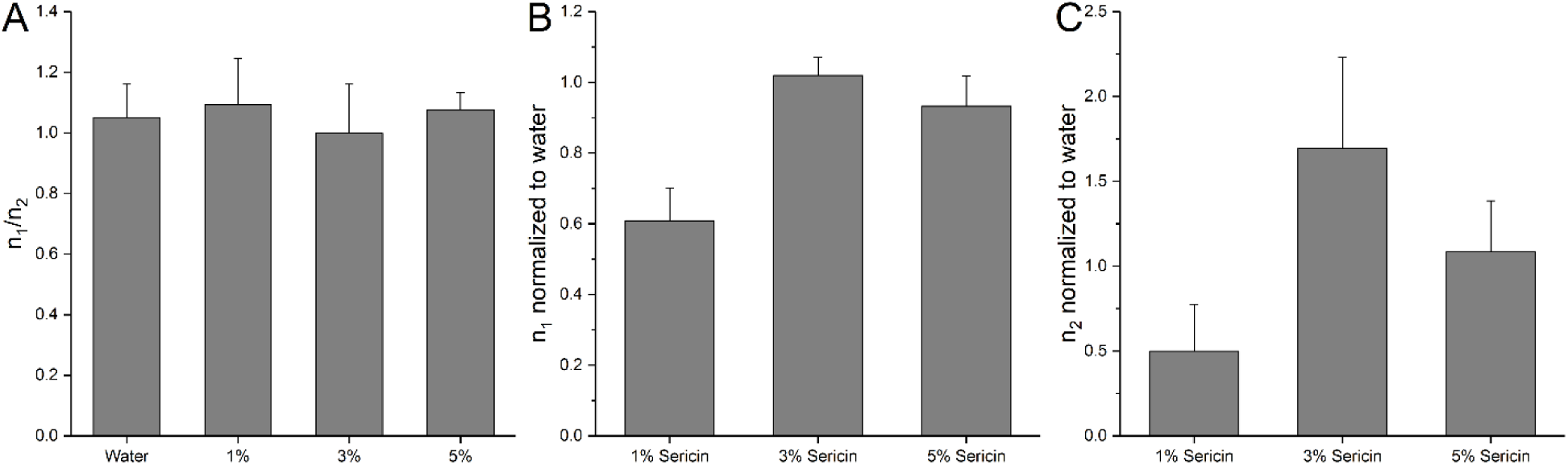
Raman OH stretching band shift hydrogen bonding analysis A) The ratio of symmetric and asymmetric slopes for water-sericin solutions. There is no statistically significant difference between the four solutions, indicating equal shift with respect to temperature for all solutions. B) The ratio of sericin solution symmetric slope over pure water slope. The normalized slope is significantly lower for the 1% solution indicating less change in hydrogen bonding as temperature is decreased. *p<0.05. C) Similar slope comparison as in (B), here for asymmetric slope. Significantly lower normalized slope for 1% sericin solution indicates lower H-bonding. Error bars show ±SEM, n=3

### Differential scanning calorimetry (DSC) for determining thermodynamic properties

The effect of addition of an additive to the CPA formulation was analyzed using DSC. Properties including freezing point, melting point, and heat of fusion was quantified using DSC. DSC measurements were performed using a precise temperature-controlled microscopy stage and a temperature controller (FDCS 600, Linkam Scientific Instruments, Tadworth, UK). Calibration of the system was performed using indium as described in ASTM E968 ([41] - Data not shown). CPA formulations containing trehalose and sericin as additives were analyzed. Thermodynamic characteristics of DMSO and sericin solutions were compared to each other (Fig. 5). The thermodynamic characteristics for 5% and 10% DMSO (v/v) in presence of additives were also compared (Fig. 6). Freezing/melting data was procured at 1°C/min until stabilized after freezing. Samples were heated at 1°C/min until stable after melting. The freezing point and melting point were considered as the temperatures at which maximum heat flow occurred during the phase change process. Enthalpy of freezing was determined by measuring the area under the curve of the thermogram. Energy data was normalized to the mass of the solution added to the DSC chamber. All thermogram data were analyzed using Origin Pro 2018.

**Figure 5:**
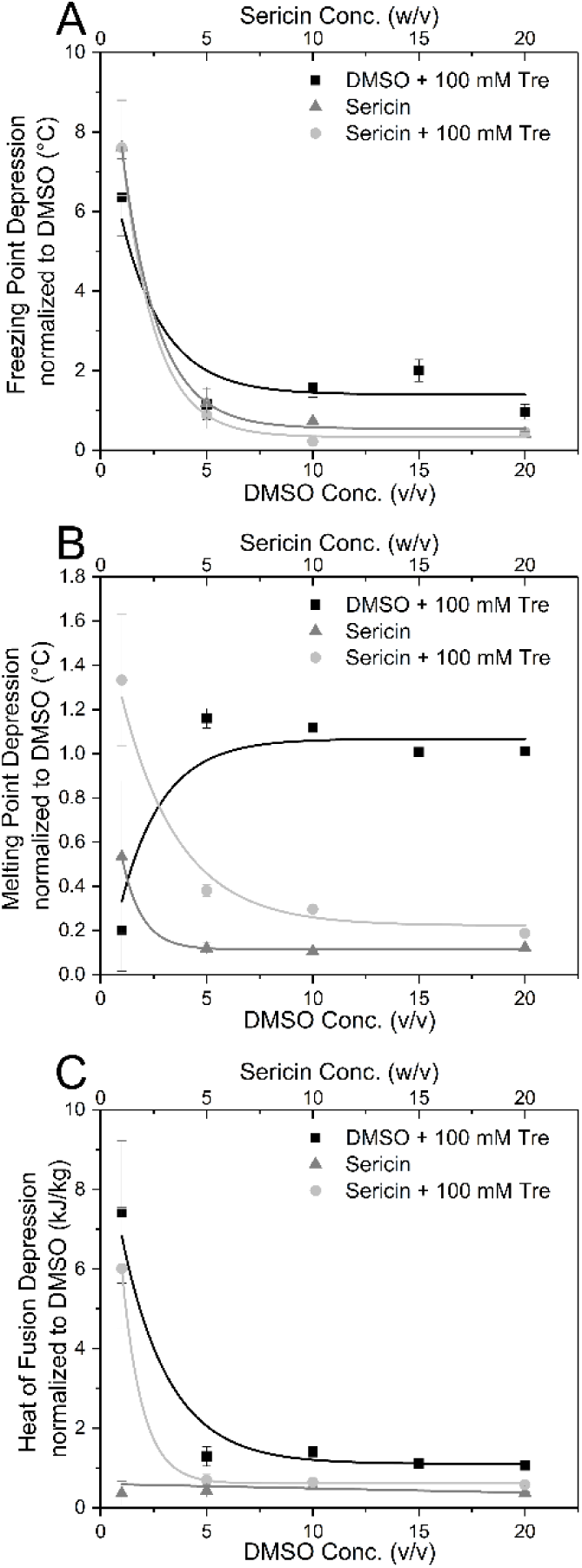
DSC analysis of individual CPA constituents A) Thermodynamic parameters were acquired for solutions with varied CPA compositions. Solutions were created with concentrations of 0-20% of DMSO (v/v) or sericin (w/v). These solutions were also analyzed with an addition of 100 mM trehalose. All parameters were normalized to DMSO solutions at equal concentrations. Freezing point depression had large immediate increases for all conditions, DMSO had more significant effects at the highest concentrations. B) All solutions had increasing melting point depression with increased CPA concentration. C) Heat of fusion depression had similar trends as melting point depression, but with DMSO and sericin solutions having closer m values. Error bars show ±SEM, n=3.

**Figure 6:**
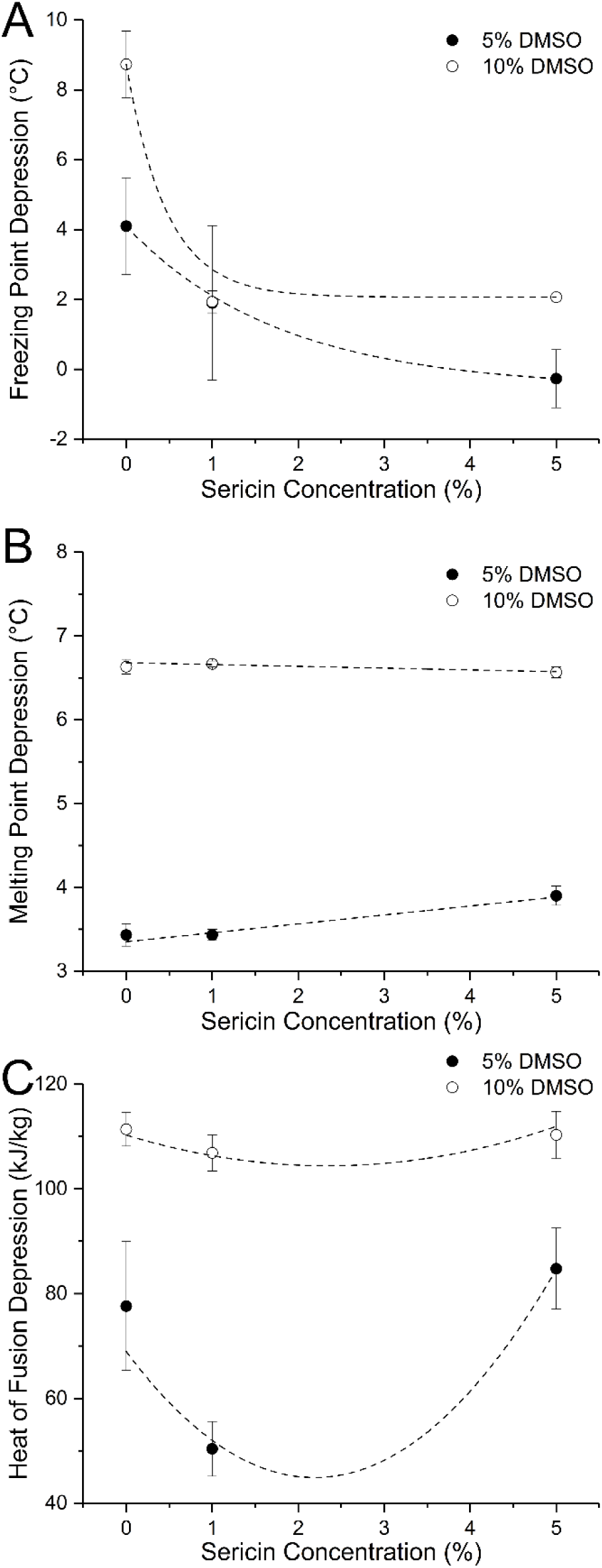
DSC analysis of CPA solutions A) CPA solutions are made with pure water and contain 100 mM trehalose, either 5% or 10% DMSO, and changing concentrations of sericin (0-5% w/v) Freezing point depression is plotted using an exponential decay function, which decreases with increasing sericin concentration (w/v). B) Change in sericin concentration has small effect on the change in melting point. C) A localized drop of heat of fusion depression at 1% sericin is highly pronounced with 5% DMSO. Error bars show ±SEM, n=3

### Cell culture, Cryopreservation and growth

Human hepatocellular carcinoma (HepG2) cells were obtained from the American Type Culture Collection (Manassas, VA), and grown in 75 cm^2^ culture flasks (Corning Inc, Corning, NY). Opti-MEM (Gibco) culture media was supplemented with 5% fetal bovine serum (FBS) (Gibco) and penicillin-streptomycin to yield final 100 units/mL penicillin G and 100 µg/mL streptomycin sulfate (Hyclone-Thermo Scientific, Logan, UT). Cells were incubated in an atmosphere of 5% CO_2_ and 95% air. The cells were collected using trypsinization followed by centrifugation and resuspended in 1 mL of cryoprotective solution in individual microtubes. A passive freezing container capable of controlling the cooling rate at 1°C/min (Cool Cell LX, 137 Biocision, Menlo Park, CA) was used to store samples in cryogenic conditions. After exposing the cells to cryogenic conditions for pre-determined duration, cells were thawed quickly in a using a 37°C water bath and re-suspended in fully complemented cell culture medium. Cell numbers were quantified using hemacytometer (Hausser Scientific, Horsham, PA) counts and membrane integrity was assessed using trypan blue exclusion. Following the initial viability count, cells were put into 25 cm^2^ flasks for the grow out. Each thawing condition had 3 separate flasks to be counted on days 3, 5, and 7. Cells were counted using the hemacytometer-trypan blue exclusion.

## Results

### Raman microspectroscopy

The effect of the presence of sericin on the H-bonding strength was investigated using Raman microspectroscopy. Fig. 1A and B shows the representative Raman spectrum of pure water at 20°C and −10°C. An FFT based chemometric algorithm was used to deconvolute the OH stretching region (∼2800-3800 rel. cm^-1^). Among the identified peaks, symmetric and asymmetric OH stretching peaks (peaks 1 and 2 respectively) were used to quantify the H-bonding characteristics in the solution state [38, 39]. The intensity variations in the deconvoluted peaks with different temperatures indicate relative changes in the nature of H-bonding with water and its neighboring molecules [37]]. Fig. 1C plots the maximum intensities of the symmetric and asymmetric peaks from −10°C to 20°C. With the increase in temperature, symmetric bonding intensity decreases linearly, while increasing the asymmetric bonding intensity. As temperature decreases, the symmetric peak intensities increase due to increase in number of central H_2_O molecules completely bound by its nearest neighbors. An opposite effect is observed for the asymmetric peak which is associated with the incompletely H-bonded clusters of water molecules in the solution.

Presence of an additive to influence the H-bonding characteristics at low temperatures can thus be quantified by comparing the pattern of increase (or decrease) in peak intensities in OH stretching regions as shown in Fig. 1C. At lower temperature this variation in overall strength of H-bonding in water clusters can be represented by the slopes of the fitted trends (m_1_ and m_2_). Fig. 1D plots the ratio of m_1_ and m_2_ for different concentrations of sericin in water (1-5% w/v). Sericin in higher concentrations is shown to decrease the contribution of incomplete water clusters and increase the overall number of strongly H-bonded water clusters at lower temperatures (Fig 1D). At higher concentration of 5% sericin, there appears to be a significant increase in rate of change in asymmetric peak intensity with temperature. In addition to the change in number of H-bonded water clusters, the functional OH groups on sericin molecules may be a contributing factor to the intensity variation in asymmetric peak intensity.

In order to understand the thermodynamic effect of presence of sericin molecules in water, a Van’t Hoff analysis was performed. Change in the enthalpy of water-sericin binary solutions relative to the enthalpy values of water was created based on Van’t Hoff plots (Fig. 2B). The change in enthalpy (Fig. 2B) and entropy (Fig. 2C) was used to quantify the change in the number of H-bonded water clusters in aqueous solutions having varying concentrations of sericin. Shift in the pattern of enthalpy change with temperature in presence of sericin can be attributed to alteration of H-bonded water clusters. With the addition of 1% sericin, a 64% decrease in enthalpy related to H-bonding. However, such dramatic depression is not observed at higher concentrations leading to significant decrease in ΔH values in the solutions containing higher concentrations of sericin. The entropy of reaction (ΔS) was formulated as a product of the y-intercept of the Van’t Hoff plot and universal gas constant. When compared with the entropy of reaction in comparison to water, a similar trend as observed with the enthalpy values is observed (Fig. 2C). Fig. 2D indicates the isokinetic relationship obtained from the ratio of the entropy and enthalpy of the sericin solutions at different temperatures.

In addition to the spectral peak intensities in the OH stretching region, the spectral shift of the symmetric and asymmetric peaks is also related to the energetics related with the in H-bonding characteristics [34, 42]. Fig. 3A and 3B indicate the peak shift patterns related to symmetric and asymmetric peaks respectively at different temperatures between 20^°^C to −10^°^C. Figures 3C and 3D show the difference in peak-shift characteristics between the temperatures 20^°^C and −10^°^C for both symmetric and asymmetric peaks. It is interesting to note that the solution containing 1% sericin presents a significantly different trend indicating significantly reduced peak-shift characteristics for both symmetric and asymmetric peaks. At higher temperatures the symmetric peaks show a minor trend of peak shift towards lower wave numbers with increase in sericin concentration (Fig. 3C). However, no such trend is observed for the asymmetric peak shift (Fig. 3D). At lower temperatures, addition of 1% sericin causes the symmetric peak to move towards higher wavenumbers, while with the increase in sericin concentration the trend is reversed. Similar trend is observed for the asymmetric peak at lower temperature indicating unique trend in H-bonding strength at 1% sericin concentration.

At high temperature, there is no change between the peak center value. This shows sericin’s ability to modulate hydrogen bonding (either stronger or weaker) as temperature is decreased toward freezing. Error bars show ±SEM, n=3

When the peak-shift characteristics for both symmetric and asymmetric peaks are compared against each-other (n_1_/n_2_), there are no appreciable difference between any of the trends compared for any of the solutions including pure water (Fig. 4A). When the slopes of the trend in peak shifts as observed in Fig. 3A and 3B were normalized against the trend in peak shift exhibited by pure water, the peak shift characteristics of 1% sericin solution appear to be significantly different for both symmetric and asymmetric peaks (Fig. 4B and 4C).

### Differential Scanning Calorimetry Studies

A comparative analysis was undertaken to evaluate thermodynamic responses of CPA formulations containing DMSO, trehalose and sericin using standard DSC techniques. Fig. 5A indicates trends in freezing point depression in CPA formulations containing 100 mM trehalose in sericin and DMSO. The trends were compared to the freezing point depression trend observed in CPA solutions containing DMSO only. These thermodynamic responses were collected at varying concentrations of DMSO and sericin. Addition of 100 mM trehalose results in significant decrease in freezing point depression characteristics when compared to DMSO-water binary solutions. Sericin-water binary solutions indicate the same initial trend, however exhibit much lower levels of freezing point depression characteristics when directly compared with DMSO-water binary system. Addition of 100 mM trehalose to sericin based solutions leads to even further reduction in trend indicating a collaborative effect of sericin and trehalose.

Fig. 5B indicates the trends in melting point depression in CPA formulations. Upon comparing the trends in melting point depression with DMSO-water binary solutions, one can observe that at 1% concentration the addition of 100 mM trehalose significantly decreases the melting point depression. Solutions with a concentration of 5% DMSO and above containing 100 mM trehalose as additive show similar melting point depression trend to DMSO-water binary solutions, however trehalose has little effect when DMSO concentration is above 15%. Both sericin based solutions have a minimal effect in melting point depression compared to DMSO solutions. Highest amount of melting point depression is achieved in the CPA formulation containing 1% sericin and 100 mM trehalose.

When the heat of fusion characteristics of the CPA formulations were compared to DMSO-water binary solution (Fig. 5C), addition of 100mM trehalose has a large impact on heat of fusion with 1% DMSO solution. However, for CPA formulations containing higher concentrations of DMSO, addition of same concentration of trehalose does not have such appreciable effect. Similar trend is observed in CPA formulation containing 1% sericin and 100 mM trehalose. CPA formulations containing sericin-water binary solutions have similar heat of fusion characteristics as DMSO-water based CPA formulations.

As a collaborative effect of trehalose and sericin was observed in DSC thermograms described above, a DSC study was undertaken with CPA formulations containing DMSO (5% and 10%) with varying amounts of sericin and 100 mM trehalose. When sericin concentration is increased from 1 – 5%, a significant increase in freezing point can be observed in CPA formulations containing both 5% and 10% DMSO (Fig. 6A). Progressive addition of sericin result in marginal but linear increase in melting point for CPA formulations containing 10% DMSO, whereas in formulations containing 5% DMSO, an opposite trend is observed (Fig. 6B). Heat of fusion values in 5% DMSO based CPA formulation show a significant reduction initially with increase in sericin concentration. However, the trend is reversed on further addition of sericin. A similar trend with lower heat of fusion values is observed in CPA formulations containing 10% DMSO (Fig. 6C).

### Membrane Integrity and Growth Patterns of Hepatocarcinoma Cells Following Cryopreservation

Fig. 7A indicates the post-thaw membrane integrity of the HepG2 cells cryo-processed using different CPA formulations. Membrane integrity for 5% and 10% DMSO-only cells were 46% and 73%, respectively. The addition of trehalose to the CPA formulation containing 10% DMSO resulted in increase in membrane integrity. Same trend in observed in CPA formulation contain 5% DMSO. However, addition of 1% sericin to CPA formulation containing 10% DMSO solutions without trehalose resulted in an increase in a 20% increase membrane integrity. No additional gain in membrane integrity is achieved by increasing the concentration of sericin. For CPA formulation containing 5% DMSO, addition of 1% sericin results in a similar increase in membrane integrity. However, addition of 100 mM trehalose to CPA formulations containing 1% and 5% sericin in 10% DMSO resulted in a decrease of membrane integrity. The same trend is observed in CPA formulations containing 5% DMSO, where maximum loss of viability (25%) is observed in solutions containing both 5% sericin and 100 mM trehalose.

**Figure 7:**
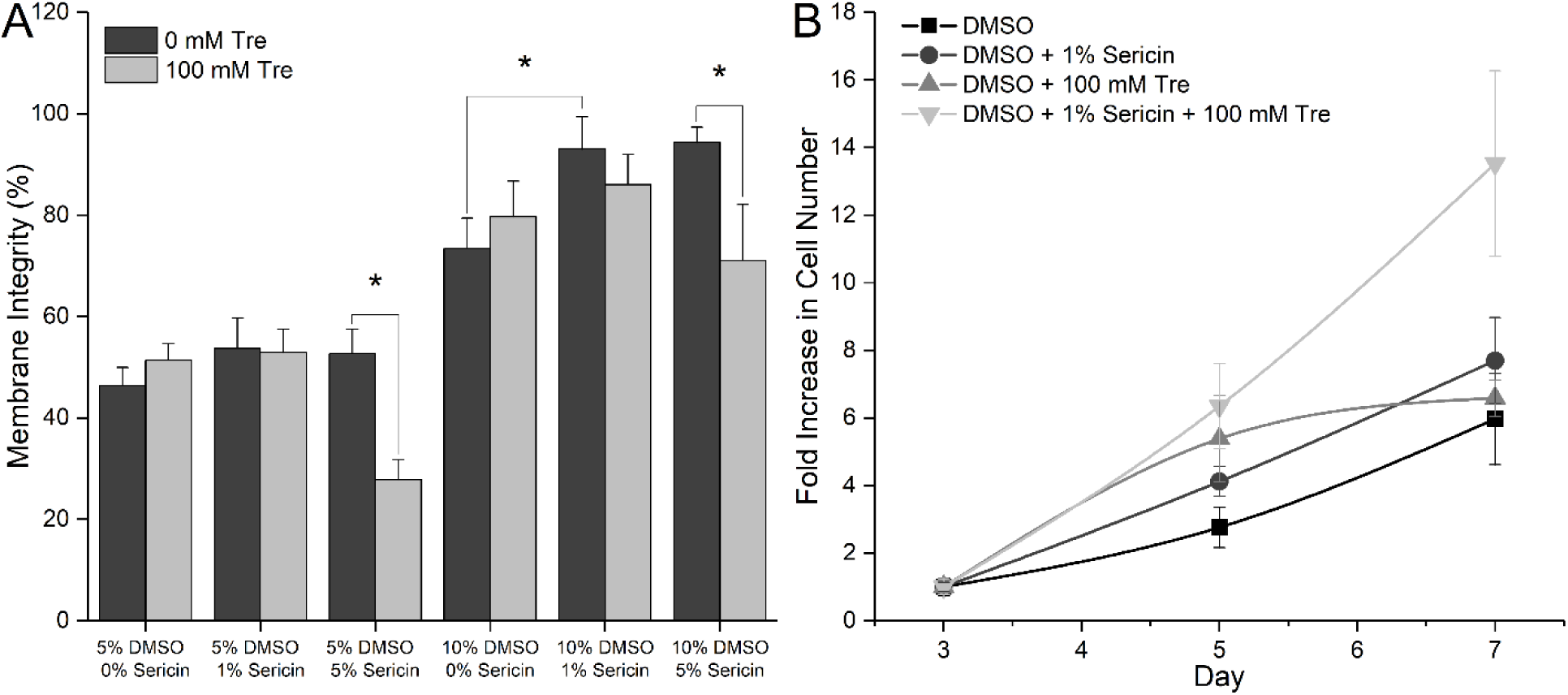
Health outcomes of mammalian cells for various CPAs A) Immediate membrane integrity for HepG2 cells after freezing for various CPA solutions with and without trehalose. Solutions containing 10% DMSO overall showed higher membrane integrity. B) After thawing, cells grown in parallel were counted for their respective conditions, all having a DMSO concentration of 10% (v/v). Data is displayed as fold increase, normalized to day 3 counts. The solution containing 10% DMSO, 1% sericin, and 100 mM trehalose had the best outcomes. Error bars show ±SEM, n=3.

When cells cryo-processed in CPA formulations containing DMSO and sericin were returned to culture conditions (Fig. 7B), the cells that survived freezing show no negative effects in cell growth (Fig. 7B). The 10% DMSO, 1% sericin, 100 mM trehalose CPA solutions showed the strongest post-thaw growth.

## Discussion

Interactions with water molecules have been shown to play a crucial role in regulating the function of biomolecules that are important to maintain cellular function [43]. Characterization of molecular level interactions is critical for unraveling the complexities of biological outcomes related to cryopreservation. Such characterization of CPA formulations in aqueous environment may hold the key to developing low toxicity cryopreservation solutions that can advance the state of the art in cryopreservation [7, 44].

CPA formulations containing 5–10% DMSO are standard in cryopreservation due to its ability to recover viable cells following freezing [4, 5, 45]. However, the major drawback of using DMSO in a CPA formulation is related to strong cytotoxic effects at physiological temperatures [7, 15, 46]. Recent studies indicate significant cytotoxic effect even at lower concentrations [6]. Efforts to mitigate cytotoxic effects of DMSO have been a major driver for using additives such as trehalose, sucrose and proline in CPA formulations [14, 18, 19, 47]. In this regard, water soluble long-chain polymers have been explored as CPA additives. Several long-chain polymers have already been explored as CPA additives due to their ability to reduce intracellular water content by increasing extracellular osmolality [48] which reduces chances of cellular injury due to IIF at low temperatures. Various serums, including fetal bovine serum (FBS), have been used as a source of long-chain proteins in CPA formulations [49]. However, the undefined nature of the majority of serums, potential presence of endotoxins, viral toxins, and other xenobiotic components make addition of serum an inherently risky and unreliable [50].

Sericin has been primarily explored as a serum replacement agent in CPA formulation. Sasaki et al. successfully used sericin as a serum replacing agent to cryopreserve human dermal fibroblasts, human epidermal keratinocytes, the rat phaeochromocytoma cell lines [29]. Human adipose tissue-derived stem cells cryopreserved in CPAs containing 1% sericin along with maltose and 10% DMSO were shown to have a superior post-thaw viability compared to those cryopreserved in similar CPAs that replaced sericin with serum [28]. However, no significant differences in the viability outcome of pancreatic islets was observed between CPA formulations containing 1% sericin and those containing 10% serum [51]. Bovine embryos cryopreserved in CPA supplemented with various concentrations (0.1%, 0.5%, and 1.0%) of sericin exhibited similar trends in survival and development to those of embryos frozen in CPA supplemented with 0.4% bovine serum albumin and 20% fetal bovine serum.

While most of these studies evaluated use of sericin to act as a serum replacement agent, the current study focuses on characterization of the possible underlying physico-chemical and thermodynamic effect of presence of sericin in CPA formulations. These effects were linked to the biological outcome and the knowledge gained here can be extended to formulate CPA formulations with superior understanding of the relationship between physico-chemical and thermodynamic parameters with post-thaw cellular viability. This can lead to the development of CPA formulations with reduced or no DMSO that has minimal potential to inflict damage in post-thaw conditions. The effect of addition of sericin in CPA formulation was evaluated using Raman microspectroscopy and differential scanning calorimetry studies.

### H-bonding Characteristics

While the role of H-bonding in modulating cryopreservation outcome is well accepted [34, 52, 53], there have been very few studies that directly link the H-bonding characteristics with the contents of CPA formulations. While some molecular dynamic simulation studies have looked at the effect of presence of CPA components such as DMSO, ethylene glycol and glycerol on H-bonding [53], experimental verification of the ability of the CPA components to influence H-bonding with water molecules is lacking. In the present study, a combined Raman microspectroscopy and DSC based analysis was undertaken to understand the ability of a sericin to influence H-bonding characteristics.

A number of techniques, including Raman [54], NMR [55], X-ray [56], neutron diffraction, and femtosecond spectroscopy have been used to study effect of water molecules to understand associated physico-chemical effect. The Raman microspectroscopy technique used in this study is an excellent tool to characterize and quantify the effect of H-bonding at molecular level in an aqueous environment at low temperatures [19, 57]. The technique was used to characterize and quantify the nature of the H-bonding network by following changes in constituent peak intensities in the OH stretching regions in the Raman spectra in presence of sericin at different temperatures. At lower temperatures the number of symmetrically arranged water clusters having lower energy forms increase in pure water causing a characteristic increase in intensity of the symmetric OH stretching peak. The opposite trend is observed for asymmetric OH stretching peak representing the incomplete water clusters. This observation is consistent with the cluster flickering phenomena described by Frank et al. and can be quantified by comparing the rate of changes in the increase of decrease of intensities of symmetric and asymmetric peaks [57, 58]. It was found that at lower temperatures, presence of sericin influences the ratio described here and at higher sericin concentrations, it decreases significantly due to decrease of incomplete water clusters and increase in overall number of strongly H-bonded water clusters. This causes a fundamental change in overall strength of H-bonding in water clusters in presence of sericin. Presence of DMSO in water-DMSO binary solutions is known to reduce the number of incomplete water clusters at lower temperature in similar fashion [59, 60]. Considering the relationship between the number of symmetrically bonded water clusters and ice crystal formation, this indicates an ability of sericin to modulate ice crystal formation. This is highly significant given the fact that the number and size of ice crystal formation have been directly linked to the post-thaw viability outcome in several studies [19] and indicates the possible role played by sericin as an additive.

The effect of presence of sericin on formation of H-bonding in water clusters were extended to quantify the thermodynamic relationships. Van’t Hoff analysis was used to quantify enthalpy of H-bond formation using the spectral data (Fig. 2B). The results indicate that the sericin can influence both enthalpy and entropy of solution by modulating H-bonding interactions with water. The strongest effects were observed at a sericin concentration of 1% w/v where a difference of over 60% was noticed compared to pure water. The significant reduction in enthalpy possibly indicates formation of extensive H-bonding network in presence of 1% sericin. The increased order of the water clusters in comparison to pure water due to increase in H-bonding will result in a decrease of both entropy and enthalpy as seen in Fig. 2B and C. However, with increase in sericin concentration, the trend is lost and a possibly indicates increase in incompletely formed water clusters and this conclusion is supported by spectroscopic observations indicated in Fig. 1D. change in as indicated by the significant decrease in both entropy and enthalpy. This is an important aspect that can be used to understand the critical parameters related to the extracellular environment at lower temperature in presence of sericin. While DMSO has a similar effect on enthalpy and H-bonding characteristics, studies indicate an absence of such trends with increasing concentrations [61, 62]. The overall linear relationship between enthalpy and entropy in aqueous solutions of sericin (Fig. 2D) in spite of significant depression at 1% w/v concentration implies a strong trend of entropy-enthalpy compensation. This compensatory trend of entropy and enthalpy is a classic indicator of involvement of H-bonding dynamics [63]. This also indicates that the change in H-bonding characteristics due to presence of sericin may also influence the degree of steric hindrance of the molecules in the aqueous environment [64]. At a concentration of 1% sericin, a coordinated decrease in entropy of activation can be achieved with the increased steric hindrance. However, it is important to note that the degree of steric hindrance caused by the nature of the H-bonding network in presence of 1% sericin follow the overall reaction framework defined by the isokinetic line. Furthermore, in such an energetic framework, the reaction of forming a symmetric bond from asymmetric bond is significantly more favorable for 1% sericin solution in comparison with solutions containing higher amount of sericin.

Along with the intensities, a shift in peak positions in OH stretching region indicates the change in energy characteristics of associated water clusters [34]. A shift towards lower wavenumbers indicate lower energy transition while a shift towards higher wavenumber is generally associated with transition to higher energy state. The dynamics related to change in wavenumber of peak positions in OH stretching region in liquid water is often connected to the reorganization of H-bonding in local solvent network [65]. However, being a collective phenomenon, it is difficult to quantify the extent of reorganization in the local H-bonding network. Recent studies indicate that a linear relationship between the change in wavenumbers of the peaks in OH stretching region and the charge and energy transfer through donor-acceptor water pairs in water-clusters. This linearity of a hydrogen bond can be related to the bond stretch frequency exhibited by its components. Increased hydrogen bonding strength shifts the OH stretching band toward lower wavenumbers [34]. This shift is noticeably observed during the formation of ice, when the spectra shifts from a broad OH peak to a sharp OH band at a much lower wavenumber [66]. Both the symmetric and asymmetric peaks were found to exhibit a shift towards low wavenumber as temperature decreases. The ratio of the amount of shift with change in temperature remains approximately proportional for all solutions tested (Fig. 4A). Interestingly, observed differences between the solutions only become significant at lower temperatures (Fig. 3C, 3D). This possibly indicates that sericin actively reduces the energy associated with the H-bonded water clusters at lower temperatures. While further studies are required to fully understand the effect of such behavior, it can be said that the rate of energy shift associated with H-bonded structure is more prominent among the water clusters incompletely bound by H-bonding (Fig. 3B). For solutions containing 1% sericin, the symmetric and asymmetric peak shifts to the right are significant compared to water, indicating an overall weakening in bonding strength at low temperatures. These results agree with the general trend observed in Van’t Hoff analysis discussed above.

### Thermodynamic analysis of freezing characteristics

Spectroscopic studies were useful in developing an understanding of the fundamental effect of sericin on H-bonding and the thermodynamic parameters derived from DSC studies provide valuable insights to relate the concentrations and compositions of CPA to the post-thaw viability. The slow cooling cryopreservation technique employed here, extracellular ice nucleation in supercooled condition is guided by the thermodynamic properties of the individual components of CPA formulation. A decrease in extracellular ice nucleation has been traditionally linked to the probability of incidence of generally lethal IIF in multiple studies [4, 48, 67]. Decreasing the nucleation temperature in extracellular environment is accompanied by an increase in the amount of intracellular supercooling; thus, separating the effects of temperature and supercooling can be challenging. Furthermore, increasing the concentrations of the non-permeating components of CPA formulations generally increase the tonicity of the extracellular solution and in turn causes the cell volume to decrease and intracellular osmolality to increase [48]. This decreases the probabilistic incidence of IIF even at higher degree of supercooling. This may be one of the contributing factors that can explain the increase in viability observed when certain additives are included in CPA formulation in addition to traditional permeating CPAs such as DMSO [18, 19]. However, there is a possibility that additives such as trehalose and sericin may have influence the thermodynamic properties of the extracellular solution in a unique way due to their mutual interaction. When 100 mM trehalose is added to sericin solutions at different concentrations, a significant loss of freezing point depression trend is observed (Fig. 5A). In addition to probabilistic decrease of IIF, such increase in freezing point may also prevent chilling injuries [68]. The collaborative effect of sericin may also have a role to play in preventing re-crystallization injury. Studies with human oocytes [67] indicate that post-thaw survival of cells can be maximized and incidence of IIF can be minimized by raising freezing point close to the melting point of CPA formulation. Sericin on its own has a similar effect on melting point compared to DMSO, and addition of trehalose decreases the melting point even further, indicating the combined effects of trehalose and sericin (Fig. 5B) may achieved better outcomes. Furthermore, the variation in heat of fusion value can be directly correlated to the difference in the nature of extracellular ice crystals formed [69]. Even though in a multi-component system, the concept of latent heat is complicated by the possible internal melting and freezing at microscale that can happen over a wide temperature range [70], a variation in heat of fusion will generally indicate a change in the overall quantity of the ice crystals formed [71]. As seen in Fig. 5C, increasing the concentration of sericin in CPA formulation on its own do not change the quantity of ice formation at the same freezing rate compared to DMSO. However, with an addition of 100 mM trehalose the heat of fusion increases indicating a significant change in the number of the ice formation. This observation is supported by detailed Raman microspectroscopic study of the nature of the ice crystal formation in presence of 100 mM trehalose in DMSO solution by Solocinski et al. [19]. Addition of 100 mM trehalose in sericin solution has a similar effect and one can assume similar trends in ice crystal formation as in a DMSO solution containing 100 mM trehalose. This is another indication of trehalose and sericin can collaboratively modulate the ice formation phenomena in extracellular environment during cryopreservation.

To further investigate the viability of sericin to be used as an additive with the capability to partially replace DMSO as a CPA component, a detailed DSC study with CPA formulations containing 5% and 10% DMSO (v/v). While 100 mM trehalose was maintained as a component, the sericin concentration was varied from 1-5% (w/v). Freezing point depression characteristics (Fig. 6A) indicate that with reduced DMSO concentration, sericin can influence the freezing point depression temperature of the CPA formulation at lower concentration. While a similar trend holds for CPA formulation containing 10% DMSO, the extent of freezing point depression is higher when compared to the solution containing lower concentration of DMSO. The difference between two CPA formulations containing different concentrations of DMSO was more prominent (Fig. 6B) when it comes to melting point depression characteristics with solutions containing 5% DMSO having a significantly lower level of melting point depression characteristics. This underscores the strong influence of DMSO on melting point depression characteristics in CPA formulations. Finally, the heat of fusion characteristics of the solutions (Fig. 6C) indicate a significantly different trend in the number of ice crystal formation. All of these indicate an increased possibility of IIF for CPA formulations containing 5% DMSO increasing the chances of cellular injury. However, with the addition of sericin and trehalose as an additive to CPA formulations with higher DMSO content, there is a chance of increased hyperosmotic exposure and related solution effects injury.

### Cellular Viability and growth

The membrane integrity of HepG2 cells cryopreserved using the CPA formulations described here indicate a higher viability of CPA formulations containing 10% DMSO (Fig. 7A). This observation is consistent with increased probability of intracellular damage due to formation of IIF when CPA formulations containing 5% DMSO as predicted by the calorimetry studies. While the addition of 100 mM trehalose to 10% DMSO solution led to an increased membrane integrity in comparison to 10% DMSO solution, addition of 1% sericin (w/v) to 10% DMSO resulted in a considerable increase in membrane integrity. Effect of addition of 1% sericin to 10% DMSO solution was higher than the CPA formulations containing 100 mM trehalose to 10% DMSO solution. This indicates the superior nature of sericin as an additive. While the thermogravimetric studies indicate distinct synergistic advantages of using both trehalose and sericin as an additive in 10% DMSO solution, the membrane integrity data indicate a 5% decrease, possibility due to increased hyperosmotic exposure and solution effects injury [48, 72]. However, it is important to note that even though the HepG2 cells that were cryopreserved using both trehalose and sericin has lower membrane integrity, their growth pattern was significantly better compared to other groups. This possibly indicate superior ability to protect the cells under post-thaw culture conditions. Further studies are underway to investigate possible protective cytoprotective effect of sericin as an additive to CPA formulation.

## Conclusion

The data presented here indicates that sericin at right concentration can play an important role as an effective additive in CPA formulations. While the usefulness of polymeric proteins in replacing DMSO in CPA formulation has been established, the study presented here provides important insight to how sericin impacts the H-bonding network and thermophysical properties of the CPA formulation during cryopreservation and provides a practical approach towards using sericin in CPA formulation as an additive to ameliorate post-thaw injuries in culture condition. While the prospect of using sericin as a replacement for DMSO in CPAs is highly attractive, further research is required to transform this study into a clinically viable CPA solution.

## Author Contributions

Conceptualization NC

Data Curation LU, JS, NC

**Formal Analysis** LU, JS, NC

**Funding Acquisition** QO NC

**Investigation** LU, ER

**Methodology** LU, ER

**Project Administration** QO, NC

**Resources** QO, NC

**Software** LU

**Supervision** QO, NC

**Visualization** LU, JS, ER, NC

**Writing – Original Draft Preparation** LU, NC

**Writing – Review & Editing** LU, JS, QO, NC

## Acknowledgements

Authors would like to acknowledge the NSF grant numbers CHE 1609440 and 1820032 Authors would also like to acknowledge instrumentation support through the startup funding received from CECS at University of Michigan-Dearborn and UM Grant Numbers U046888. NC has a financial interest in Otzi Bio LLC, a company focused on developing biopreservation technology. NC interests are managed by the University of Michigan in accordance with their conflict-of-interest policies.

